# Induction of *Bdnf* from promoter I following electroconvulsive seizures contributes to structural plasticity in neurons of the piriform cortex

**DOI:** 10.1101/2021.03.04.433962

**Authors:** Anthony D. Ramnauth, Kristen R. Maynard, Alisha S. Kardian, BaDoi Phan, Madhavi Tippani, Sumita Rajpurohit, John W. Hobbs, Stephanie Cerceo Page, Andrew E. Jaffe, Keri Martinowich

**Author notes:** Equal Contribution (Anthony D. Ramnauth, Kristen R. Maynard). Correspondence: Kristen Maynard, Lieber Institute for Brain Development, 855 North Wolfe Street, Suite 300, Baltimore, MD, 21205.

## Abstract

The efficacy of electroconvulsive therapy (ECT) as a treatment for psychiatric disorders, including major depressive disorder (MDD) is hypothesized to depend on induction of molecular and cellular events that trigger structural plasticity in neurons. Electroconvulsive seizures (ECS) in animal models can help to inform our understanding of how electroconvulsive therapy (ECT) impacts the brain. ECS induces structural plasticity in neuronal dendrites in many brain regions, including the piriform cortex, a highly epileptogenic region that has also been implicated in depression. ECS-induced structural plasticity is associated with differential expression of unique isoforms encoding the neurotrophin, brain-derived neurotrophic factor (BDNF), but the functional significance of these transcripts in dendritic plasticity is not clear. Here, we demonstrate that different *Bdnf* isoforms are expressed non-stochastically across neurons of the piriform cortex following ECS. Specifically, cells expressing *Bdnf* exon 1-containing transcripts show a unique spatial recruitment pattern in response to ECS. We further demonstrate that *Bdnf* Ex1 expression in these cells is necessary for ECS-induced dendritic spine plasticity.

## INTRODUCTION

Electroconvulsive therapy (ECT) remains one of the most effective treatments for major depressive disorder (MDD), particularly for patients living with treatment resistant depression (Kho *et al*, 2003; Pagnin *et al*, 2004; UK ECT Review Group, 2003). However, despite its use as a treatment for over 80 years, the neurobiological mechanisms by which ECT improves symptoms of depression remain unclear. Some clinical evidence suggests that ECT normalizes hypothalamic-pituitary-adrenal (HPA) axis function in depression (McKay and Zakzanis, 2010; Yuuki *et al*, 2005), and in rodent models of chronic stress exposure HPA axis dysregulation increases levels of circulating glucocorticoids, which induces dendritic atrophy (Alfarez *et al*, 2009; Goldwater *et al*, 2009; Morales-Medina *et al*, 2009). Exposure to chronic stress also decreases synaptic spine density (Duman and Duman, 2015) and alters dendritic spine morphology (Duman and Duman, 2015; Radley *et al*, 2008) in limbic brain areas that are implicated in MDD (Radley *et al*, 2008). The effects of ECT can be studied in rodent models using electroconvulsive seizures (ECS), administration of which reverses chronic stress-induced changes in neuronal and dendritic spine morphology (Chang *et al*, 2018; Kaastrup Müller *et al*, 2015; Maynard *et al*, 2018; Schloesser *et al*, 2015). In addition to reversing these stress-induced morphological changes, ECS also attenuates behaviors observed in rodent models of chronic stress exposure that are relevant for depression (Azis *et al*, 2019; Chang *et al*, 2018; Schloesser *et al*, 2015).

Decreased brain-derived neurotrophic factor (BDNF) expression is documented in multiple brain regions in rodent models of chronic stress exposure (Duman and Monteggia, 2006; Martinowich *et al*, 2007; Smith *et al*, 1995). Additional studies have correlated BDNF expression increases with effects of pharmacological antidepressant administration as well as ECS (Autry *et al*, 2011; Nibuya *et al*, 1995; Russo-Neustadt *et al*, 2000)). Consistent with these findings, BDNF levels are lower in postmortem human brain tissue from donors diagnosed with MDD compared to neurotypical donors (Dwivedi *et al*, 2003; Guilloux *et al*, 2012). This data, together with the fact that multiple therapies for depression increase BDNF levels in the brain (Chen *et al*, 2001; Guilloux *et al*, 2012; Kraus *et al*, 2019; Lee and Kim, 2010; Schmidt *et al*, 2011), contributed to the hypothesis that BDNF levels are a biomarker for antidepressant efficacy. In rodents, *Bdnf* transcription is controlled by 9 unique promoters, which generate at least 20 different *Bdnf* transcript isoforms, all of which encode the same BDNF protein (Aid *et al*, 2007). Each isoform consists of a 5’ untranslated region (UTR) exon, which is alternatively spliced to a downstream common coding exon (exon 9) (Aid *et al*, 2007). These isoforms are differentially expressed across brain regions as well as during neurodevelopment (Aid *et al*, 2007; Sathanoori *et al*, 2004; Timmusk *et al*, 1993). The exon 1- and exon 4-containing *Bdnf* variants (Ex1 and Ex4), which are generated from transcriptional activity at *Bdnf* promoter I and IV, respectively, are strongly induced by neural activity (Aid *et al*, 2007; Sathanoori *et al*, 2004; Timmusk *et al*, 1993). These activity-dependent isoforms also exhibit differential subcellular localization e.g., *Bdnf* Ex1 and Ex4 transcripts are localized to the cell body and proximal dendrites, while exon 2 (Ex2)- and exon 6 (Ex6)-containing transcripts are localized in distal dendrites (Baj *et al*, 2011; Chiaruttini *et al*, 2008; Pattabiraman *et al*, 2005). We previously demonstrated that both acute and repeated ECS upregulate activity-dependent *Bdnf* Ex1- and Ex4 isoforms (Maynard *et al*, 2018, 2020). We also showed that activated BDNF-expressing cells may preferentially utilize one of these activity-dependent isoforms over the other, contributing to the notion that *Bdnf* Ex1 and Ex4 expression may partially define sets of spatially and functionally discrete activity-induced neuronal ensembles (Maynard *et al*, 2020). This is important since BDNF production from different *Bdnf* transcripts has unique effects on neural plasticity and cell function (Hallock *et al*, 2019; Hill *et al*, 2016; Maynard *et al*, 2016, 2017; Savell *et al*, 2019).

Following ECS, *Bdnf* Ex1 transcripts are strongly induced in the piriform cortex (Maynard *et al*, 2020), a highly epileptogenic brain region that may also be important in depression (Ekstrand *et al*, 2001; Kohli *et al*, 2016; Soudry *et al*, 2011). To better understand the regulation of activity-induced *Bdnf* expression, we investigated spatial relationships between *Bdnf* Ex1 and Ex4-expressing cells in the piriform cortex following ECS. Identification of a unique pattern of spatial recruitment of *Bdnf* Ex1 cells prompted us to further investigate whether *Bdnf* Ex1 isoforms are required for ECS-induced dendritic spine morphology. To genetically tag *Bdnf* Ex1-expressing cells for labeling and morphological analysis, we used previously generated mice that report GFP under control of *Bdnf* promoter I in combination with viral delivery of a GFP-dependent split Cre-recombinase system (Maynard *et al*, 2016; Tang *et al*, 2015). This strategy allowed us to assess the necessity of *Bdnf* Ex-1 transcript expression for ECS-induced changes in spine density and morphology in BDNF-expressing cells within the piriform cortex. Together, these results demonstrate that cells expressing individual *Bdnf* splice isoforms have unique spatial recruitment patterns following ECS, and that *Bdnf* Ex1 transcripts contribute to structural plasticity in the piriform cortex following ECS.

## METHODS

### Animals

We used Bdnf-e1 mice, which are engineered to disrupt BDNF production from promoter I, backcrossed to C57BL/6J > 12x (Maynard *et al*, 2016). In Bdnf-e1 mice an enhanced green fluorescent protein (eGFP)-STOP cassette is inserted upstream of the 5’UTR splice donor site of exon 1 such that transcription is initiated from promoter I to generate a 5’UTR-eGFP-STOP-Bdnf IX transcript, which produces GFP in lieu of BDNF. Adult male Bdnf-e1 −/+ and Bdnf-e1 −/− mice were used for all experiments. Mice were housed in a temperature-controlled environment with a 12:12 light/dark cycle and ad libitum access to food and water. After weaning, mice were housed in divider caging at 5 weeks of age due to excessive aggression in Bdnf-e1 male animals (Maynard *et al*, 2016). All experimental animal procedures were approved by the Sobran Biosciences Institutional Animal Care and Use Committee.

### Viral Vectors and Stereotaxic Surgery

Adeno-associated viruses (AAVs): AAV2-CAG-FLEX-tdTomato (Cat #Av-1-PV2812, Penn Vector Core), AAV2-CMV-LacZ (Cell Biolabs, Catalog number AAV-342), AAV1/2-EF1a-N-CretrcintG (Cat #B387Harvard Medical School Gene Vector Core), AAV1/2-EF1a-C-CreintG (Cat #B386, Harvard Medical School Gene Vector Core). Stereotaxic surgeries were performed on animals at 10 weeks of age. 800 nL of virus was injected bilaterally into the anterior and posterior piriform cortex at 150 nl/minute using a 10 μL Hamilton syringe targeted to AP −0.34, ML +/−3.13, DV −5 (from bregma and AP −1.7, ML +/−3.5, DV −5 from bregma). Mice were administered 100 μL of 0.5 mg/mL meloxicam injected intraperitoneally and 100 μL of 5 mg/mL bupivacaine at the incision site. Staples were removed after 2 weeks. Viruses were given 2 weeks to express before initiating further experiments.

### Electroconvulsive seizures (ECS) and brain tissue collection

Mice were administered either Sham or electroconvulsive seizures (ECS) as previously described (Maynard *et al*, 2018). Mice were anesthetized using inhaled isoflurane prior to and during treatment. ECS treatment consisted of either one ECS session (acute ECS) or 7 ECS sessions across a 15 d period (repeated ECS). ECS were delivered with an Ugo Basile pulse generator (model #57800-001, shock parameters: 100 pulse/s frequency, 3 ms pulse width, 1 s shock duration and 50 mA current) (Maynard *et al*, 2018; Schloesser *et al*, 2015). Seizures were confirmed by visualization of tonic-clonic convulsion. 48 h after the final treatment session, animals were anaesthetized with isoflurane and perfused transcardially with 4% paraformaldehyde. Brains were post-fixed for 24 h and transferred to 30% sucrose. Cryosections were cut coronally at 50 μm, mounted in consecutive order onto Superfrost Plus (VWR, 48311-703) glass slides and coverslipped using Fluoromount (VWR, 0100-01).

### Confocal Imaging

Higher-order apical dendrites of TdTomato-positive pyramidal neurons in the piriform cortex were imaged in *z*-series at 63x magnification with 3x digital zoom using a Zeiss LSM 780 microscope (Carl Zeiss, Oberkochen, Germany) with the experimenter blinded to genotype and treatment. Pyramidal neurons were selected for imaging based on the following criteria: (1) adequate brightness and isolation to allow for clear reconstruction of spine density and morphology, (2) clear attachment of dendrite to cell body to allow for unambiguous identification of branch order (i.e., we did not image “floating” branches in which cell bodies were in another plane or different section), and (3) close proximity to the coverslip to allow for capture of all branches in the z-dimension. We imaged 44 dendrites from 4 brains for Bdnf-e1 −/− ECS, 35 dendrites from 4 brains for Bdnf-e1 −/− Sham, 52 dendrites from 4 brains for Bdnf-e1 −/+ ECS, and 58 dendrites from 4 brains for Bdnf-e1 −/+ Sham. Each dendrite was imaged from a different neuron.

### Dendrite Reconstruction

Individual dendrites were manually traced from confocal z-stacks using Neurolucida 360 (MicroBrightField Biosciences, Williston, VT, USA) with the experimenter blind to genotype and treatment. For quantification of spine density (spines per micrometer) and morphology, Neurolucida 360 Explorer (MicroBrightField Biosciences) was used to calculate the number, height, and width of spines on segments of dendrite branches. Student’s *t* tests were used to analyze spine density, and ordinary two-way ANOVAs followed by Bonferroni post hoc tests were used to examine differences between the percentage of differently sized spines on Bdnf-e1 +/− and Bdnf-e1 −/− branches between ECS and Sham treatments.

### Spatial distribution of cells expressing *Bdnf* and *Arc* transcripts

We applied the centered, weighted Besag’s transformation of Ripley’s K function (akin to the centered L-function) to determine the relative distribution of nuclei weighted by transcript counts. Briefly, this allows us to ask whether nuclei with more transcript counts tend to be spaced at a particular distance away from each other compared to the given spacing of the present nuclei unweighted by *Bdnf* transcript counts. We also took a complementary approach to continuous transcript counts by binning nuclei into high-, medium, and low-expressors of *Bdnf* transcripts. Subsetting to just medium and high *Bdnf* expression categories, we applied the standard L-function to ask whether med/high-expressing nuclei are spaced at a particular distance away from each other, ignoring low-expressors, and above a random spacing. Expression categories defined as previously described (Maynard *et al*, 2020) with *Bdnf* Ex1 expression based on number of transcripts (low: <11; medium: 11-25; high: >25) and average transcript size (low: <17; medium: <74; high: >0), and *Bdnf* Ex4 number of transcripts (low: <11; medium: 11-25; high: >25) and average transcript size (low: <21; medium: <59; high: >0 ()). Calculations were performed using the spatstat packing in R with the *kmark* function, which estimates the mark-weighted K function of a marked point pattern and performs the Besag’s transform to the L function (or by using *Lest* to return the standard L function), and the marks were computed by the *markcorr* function on the density of puncta between pairs of nuclei. R documentation for this package can be found here: https://cran.r-project.org/web/packages/spatstat/spatstat.pdf

## RESULTS

### *Bdnf* Ex1-expressing neurons in the piriform cortex show a unique pattern of spatial recruitment following ECS

Using single-molecule fluorescence in situ hybridization (smFISH), we previously demonstrated that activity-regulated genes (ARGs), including *Bdnf* Ex1, *Bdnf* Ex4 and *Arc* transcripts are robustly elevated in the mouse piriform cortex following a single session of ECS (Maynard *et al*, 2020). In that study, we grouped *Bdnf* Ex1, *Bdnf* Ex4, and *Arc*-expressing cells into categories of “high”, “medium”, or “low” transcript expressors (Maynard *et al*, 2020). Here, we further analyzed that dataset, assessing patterns of recruitment, and defining spatial relationships between “high”, “medium”, and “low” *Bdnf* Ex1, Ex4, and *Arc*-expressing cells in response to ECS. Visual inspection of data that plotted the spatial distribution of “high”, “medium”, and “low” *Bdnf* Ex1, Ex4, and *Arc* expressing cells suggested that these groups of cells may have divergent patterns of spatial recruitment in response to ECS (Fig. 1B). To quantify the extent to which ECS impacts spatial relationships between cells expressing these transcripts, we computed the Besag’s transformation of Ripley’s K-function (L-function). This summary statistic measures the degree of cell nuclei clustering, where positive values indicate greater difference from chance, and negative values indicate a negative difference from chance of finding pairs of nuclei which have a given distance (r) from each other; with a lower value suggesting more clustered nuclei. Following a single ECS session, all *Bdnf* Ex1-expressing cells had a lower L function than Sham treated mice as the distances increased (Fig. 1C), suggesting that ECS recruits *Bdnf* Ex1-expressing cells that are closer together. Interestingly, if cells with low levels of *Bdnf* Ex1 transcripts (< 6 puncta) are excluded, the distribution of nuclei only modestly decreases after ECS (Fig. 1F). This change in spatial distribution is unique to *Bdnf* Ex1 isoforms, since Ex4-expressing nuclei only show a modest decrease in spatial distribution following ECS (Fig. 1D). Furthermore, the changes to spatial distribution are distinct for other ARGs, as *Arc*-expressing nuclei show a different change in distribution compared to *Bdnf*-expressing nuclei with acute ECS compared to Sham (Fig. 1E, H).

**Figure 1:**
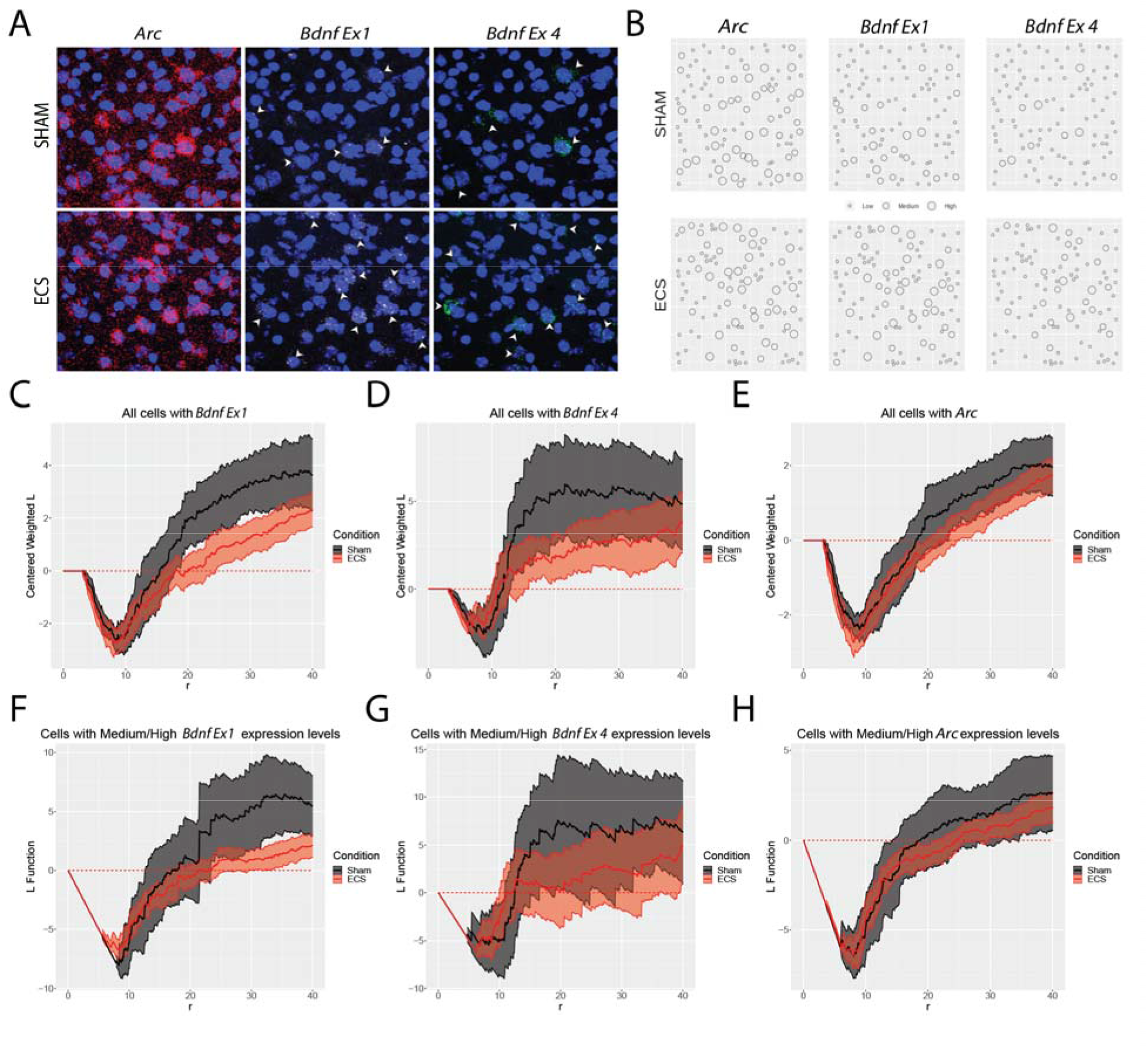
Differential spatial clustering of *Bdnf*-expressing cells in piriform cortex in response to ECS. **ECS. A** Representative images depicting expression of *Arc* (red), *Bdnf* Ex1 (white), and *Bdnf* Ex4 (green) transcripts in the piriform cortex of mice administered Sham or acute ECS; Scale bar = 20 um. **B** Point plot of representative images for nuclei. Circle diameter size represents low, medium, or high expression of indicated transcript in that nucleus as determined by k-means clustering. **C-E** Centered weighted Besag’s transformation of Ripley’s K-function (L-function) vs. radius from nuclei of all cells. Shaded areas are confidence intervals. **F-H** L-function vs. radius from nuclei of cells in medium & high k-means clusters. Shaded areas are confidence intervals.

### Selective labeling of *Bdnf* Ex1-expressing neurons using Cre recombinase dependent on GFP technology

Our previous studies showed that ECS impacts dendritic spine morphology and increases *Bdnf* Ex1 expression (Maynard *et al*, 2018). This finding, together with the unique spatial recruitment of *Bdnf* Ex1-expressing cells observed following ECS (Fig. 1), prompted the hypothesis that *Bdnf* Ex1 transcription is required for ECS-induced structural plasticity in BDNF-expressing neurons of the piriform cortex. To achieve robust labeling of *Bdnf* Ex1-expressing cells that would facilitate visualization of dendritic structures, we employed the Cre recombinase dependent on GFP (CRE-DOG) technology (Tang *et al*, 2015) in combination with a previously generated mouse line that reports GFP expression from *Bdnf* promoter I (Bdnf-e1) (Maynard *et al*, 2016). In these mice, an eGFP-STOP cassette inserted upstream of the 5’UTR splice donor site of exon 1 leads to GFP translation in lieu of BDNF (Fig. 2A). While GFP is selectively expressed in cells where there is active transcription from *Bdnf* promoter I, the fluorescent GFP signal is too weak to robustly label membrane structures such as dendritic spines for morphological analysis (Maynard *et al*, 2016). The CRE-DOG system combines two viral constructs that each encode one half of a split Cre recombinase fused with a GFP binding protein (GBP) (AAV1/2-EF1a-N-CretrcintG & AAV1/2-EF1a-C-CreintG) (Tang *et al*, 2015). We co-injected these vectors with a Cre-dependent TdTomato vector (AAV2-CAG-FLEX-tdTomato) to fluorescently label GFP-expressing cells in piriform cortex of Bdnf-e1 mice. Under these conditions, in a cell expressing GFP in lieu of *Bdnf* Ex1, the two GBP-split Cre fusion proteins complex with the GFP, which functions as a scaffold for expression of Cre-recombinase, which subsequently flips and excises the Cre-dependent tdTomato. (Fig. 2B-C). To demonstrate specificity, we executed two controls. First, when only one half of the split-Cre system (N or C terminus) is injected, TdTomato labeling is absent (Fig. 2D), and second, when CRE-DOG viruses are injected into wild-type mice along with a Cre-dependent tdTomato reporter and a constitutively expressed LacZ reporter, beta-gal staining is observed without tdTomato labeling (Fig. 2F-F”). Together, these results validate the utility of CRE-DOG technology in Bdnf-e1 mice to selectively and robustly label the subpopulation of cells in the piriform cortex in which transcription is occurring at *Bdnf* promoter I.

**Figure 2:**
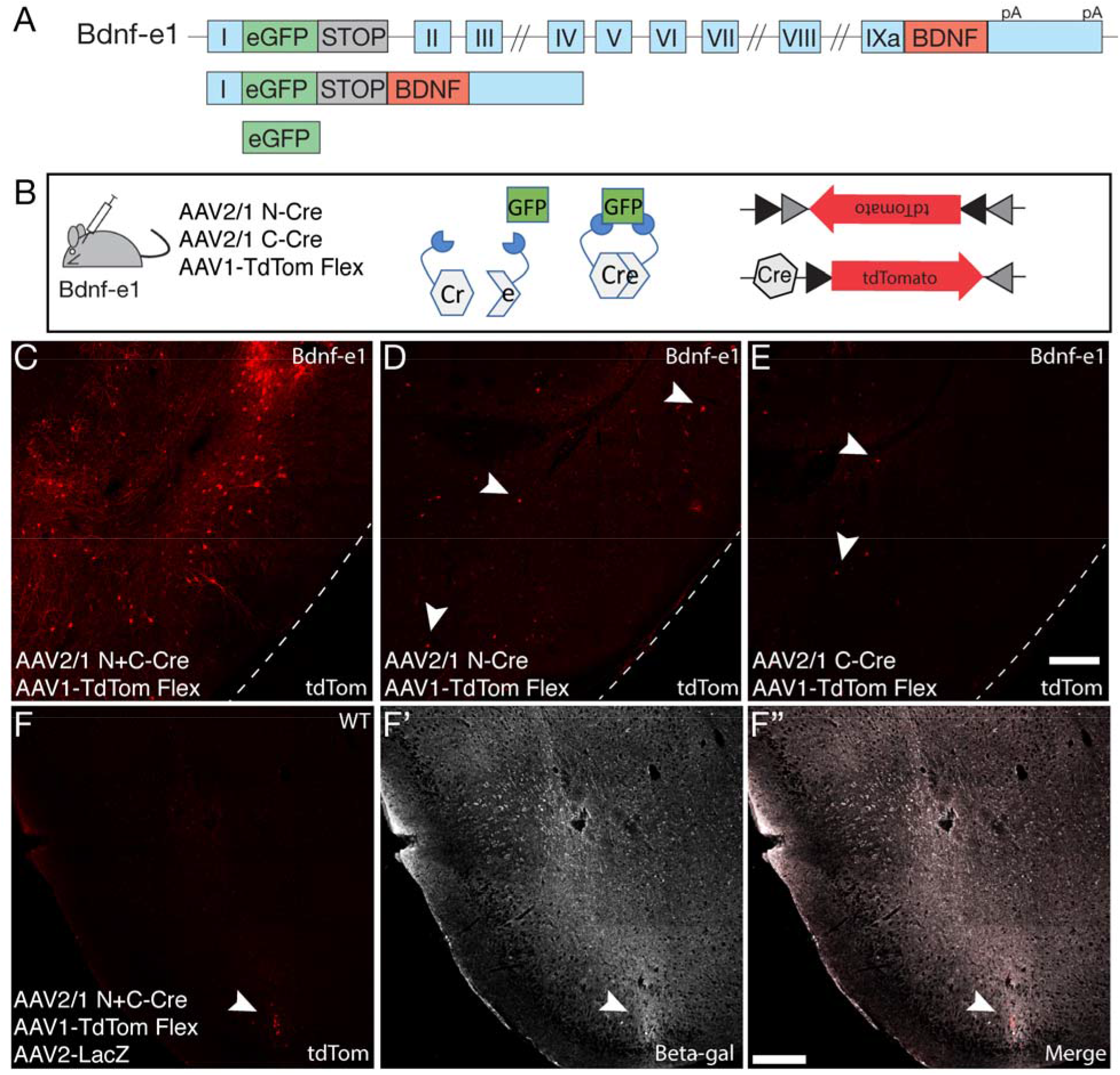
Selective targeting of *Bdnf* exon 1-expressing neurons in piriform cortex using Cre-dog technology. **A** Genetic targeting strategy for Bdnf-e1 mice in which green fluorescent protein (GFP) is produced in lieu of BDNF from promoter I **B** Function of CRE-DOG construct in Bdnf-e1 mice. CRE-DOG is a split Cre recombinase delivered to the mouse in two separate viruses that scaffold onto GFP. Expression of Cre-dependent TdTomato occurs only in GFP-expressing cells. **C** Expression of TdTomato fluorescent protein in the piriform cortex is present when N and C cre are delivered together. **D-E** Expression of TdTomato is significantly reduced when only N (d) or only C (e) CRE-DOG constructs are delivered alone **F-F”** Expression of TdTomato is absent in wild-type mice compared to Bdnf-e1 mice given the absence of GFP in wild-type mice. Beta galactosidase staining for a co-injected control LacZ constructs confirms successful viral delivery.

### Loss of promoter I-derived BDNF impacts ECS-induced spine density and morphology in *Bdnf* Ex1-expressing neurons

We next investigated whether *Bdnf* Ex1 transcripts are required for structural plasticity in dendritic spines. Using the CRE-DOG strategy with Bdnf-e1 mice, we labeled neurons reporting *Bdnf* Ex1 expression in heterozygous (+/−; intact BDNF production from *Bdnf* promoter I at one allele) and mutant mice (−/−; BDNF production from *Bdnf* promoter I blocked at both alleles). We note that in the mutant animals, the fluorescently labeled cell population is not actually expressing *Bdnf* Ex1 - these cells are only expressing GFP as a read-out of transcriptional activity at *Bdnf* promoter I. Two weeks following stereotaxic viral injection of CRE-DOG and AAV2-CAG-FLEX-tdTomato vectors into the piriform cortex to fluorescently label and fill *Bdnf* Ex1 cells, mice were administered Sham or repeated ECS (7 ECS sessions across 15d). Mice were perfused and brains collected 48 h after the final Sham or ECS session. Brains were then sectioned, and labeled neurons in the piriform cortex were imaged to quantify changes in dendritic spine morphology across these four groups: Bdnf-e1 +/−; Sham, Bdnf-e1 +/−; ECS, Bdnf-e1 −/−; Sham, Bdnf-e1 −/−; ECS. Spine density was significantly decreased in piriform cortex neurons from Bdnf-e1 −/−; ECS compared to both Bdnf-e1 −/+; Sham and Bdnf-e1 −/+; ECS mice, highlighting the importance of promoter I-derived BDNF following ECS-induced neuronal activity in dendritic spine dynamics (Fig. 3B). No change in spine length was detected between Sham and ECS for either genotype (Fig. 4A), but a significant decrease in the percentage of smaller spines (diameter 0.2-0.4 μm) in Bdnf-e1 −/− ECS mice compared to Bdnf-e1 −/+ Sham and Bdnf-e1;ECS mice was observed (Fig. 4B). These findings suggest that promoter I-derived BDNF is necessary for the maintenance of dendritic spines in *Bdnf* Ex1-expressing neurons, particularly following repeated induction of neuronal activity.

**Figure 3:**
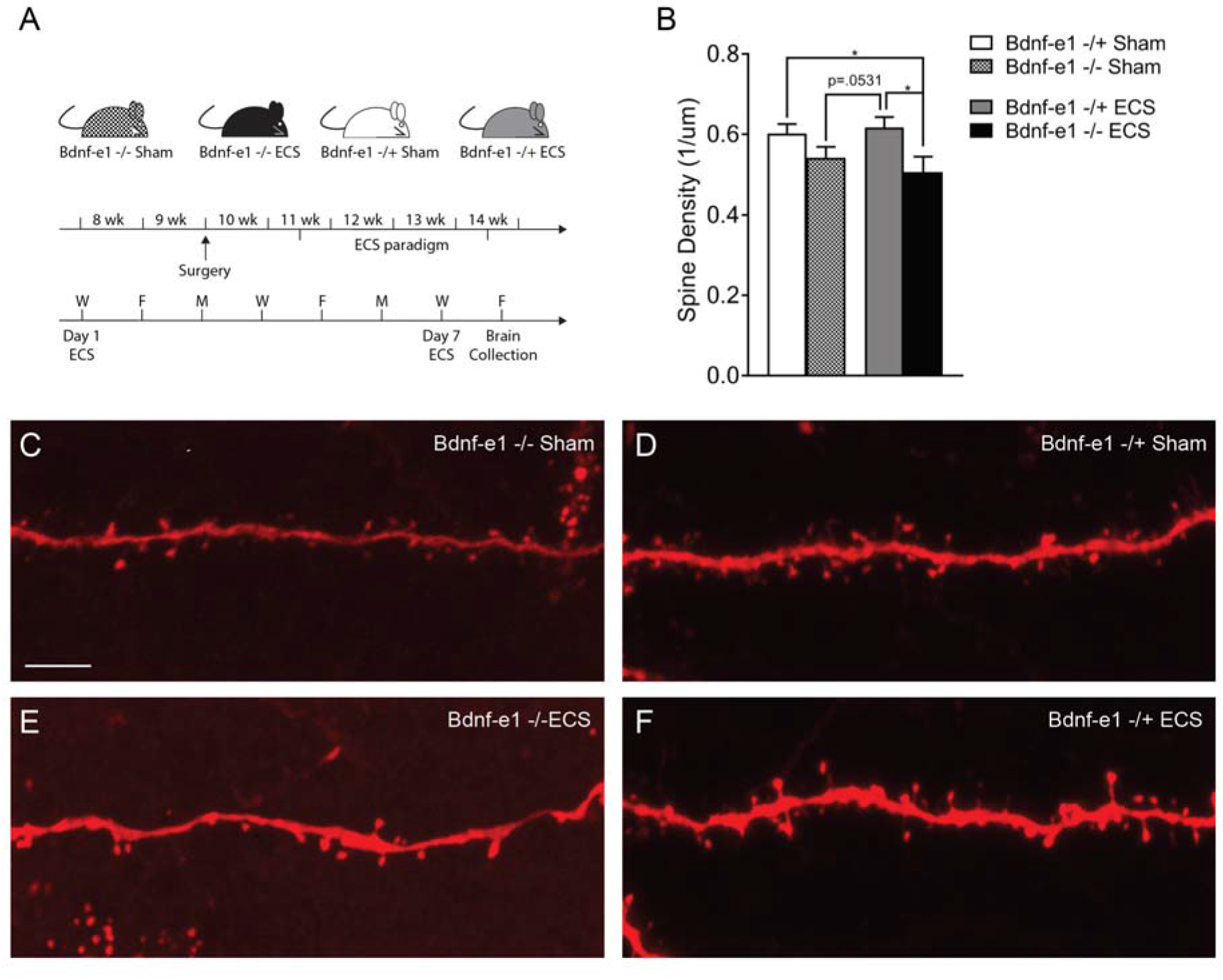
Loss of activity-induced promoter I-derived BDNF following ECS leads to reductions in dendritic spine density in *Bdnf* exon 1-expressing neurons. **A** Schematic of experimental timeline in which surgery was performed at 10 weeks followed by administration of 7 sessions of Sham or ECS treatment over 2 weeks. Brains were collected 48 hours after the final treatment session. **B** Comparison of spine density between groups. Bdnf-e1 −/− mice show lower spine density compared to Bdnf +/− mice receiving Sham or ECS treatment. For each condition n = 4 mice, 6-18 branches per mouse, 35-55 total branches per condition. Student’s *t* test, data presented as mean ± SEM *p<0.05 **C-F** Representative confocal images of dendrites and spines for each group. Scale bar is 5 um.

**Figure 4:**
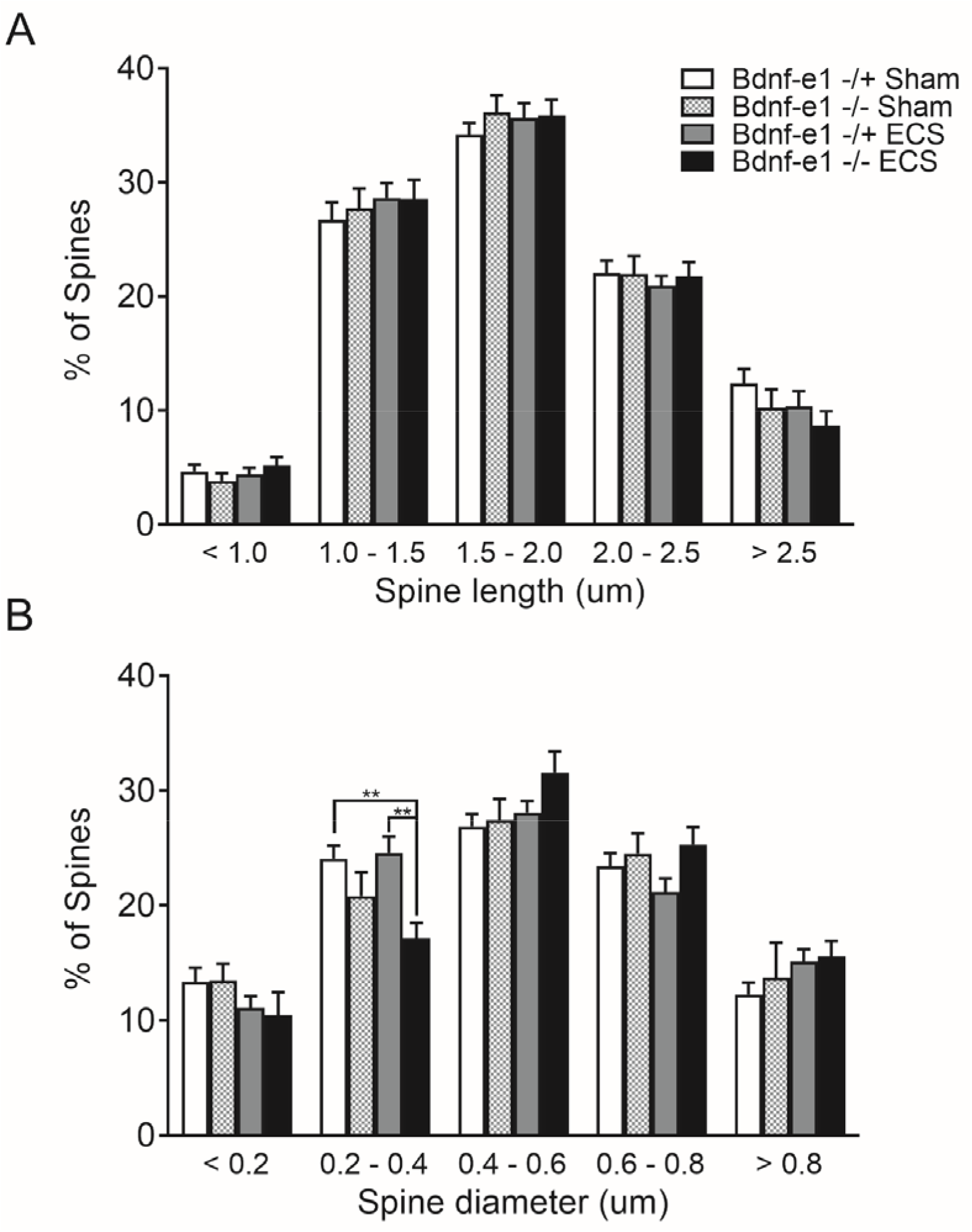
Altered spine morphology in *Bdnf* Ex expressing neurons following loss of ECS-induced promoter I-derived BDNF. **A** Neither genotype nor treatment has an effect on spine length. **B** ECS treatment causes increased number of 0.2-0.4 um diameter spines in Bdnf-e1 −/+ mice compared to Bdnf-e1 −/− mice. For each condition n = 4 mice, 6-18 branches per mouse, 35-55 total branches per condition. Data are analyzed using ordinary two-way ANOVA followed by Bonferroni post hoc tests and presented as mean ± SEM **p<0.01

## DISCUSSION

BDNF signaling is strongly linked to changes in spine density and morphology (von Bohlen Und Halbach and von Bohlen Und Halbach, 2018; Song *et al*, 2017), which are known structural changes that can functionally impact synaptic plasticity (Arellano *et al*, 2007; Hering and Sheng, 2001). These BDNF-dependent signaling events have been hypothesized to contribute to mechanisms by which antidepressant therapies relieve symptoms of depression (Autry and Monteggia, 2012). Previous studies have shown that individual *Bdnf* isoforms are differentially associated with unique BDNF-dependent molecular, cellular and behavioral functions (Hallock *et al*, 2019; Maynard *et al*, 2016, 2017, 2018; Savell *et al*, 2019). For example, at the cellular level, depleting *Bdnf* Ex1 and Ex4 isoforms in cultured hippocampal neurons reduces the number of proximal neuronal dendrites, while depleting *Bdnf* Ex2 and Ex6 isoforms decreases the number of distal neuronal dendrites (Baj *et al*, 2011). Constitutive disruption of BDNF production from specific isoforms using *in vivo* transgenic mouse models also causes divergent effects on dendritic morphology in the mouse hippocampus (Maynard *et al*, 2017). In addition, disrupting BDNF from distinct promoters results in divergent behavioral and circuit phenotypes in mice; specifically, disruption of BDNF production from promoter I results in hyperphagia and increased aggression (Maynard *et al*, 2016; McAllan *et al*, 2018), while BDNF disruption from promoter IV results in increased fear expression that is associated with disruptions in hippocampal-prefrontal cortical connectivity (Hallock *et al*, 2019, 2020; Hill *et al*, 2016). The data reported here demonstrate that the cell population expressing *Bdnf* Ex1 exhibits a distinct spatial pattern of activation in response to ECS. Specifically, recruited *Bdnf* Ex1-expressing cells are more tightly clustered together, and show higher expression of *Bdnf* Ex1 transcript per individual cell. The functional significance of local neuronal networks expressing unique patterns of *Bdnf* splice variants is not entirely clear, but highlights a preferred recruitment of *Bdnf* Ex1-expressing cells in response to ECS. Given the differential effects of BDNF secretion and signaling in the pre-versus post-synaptic compartment and consequences for impacting local autocrine and paracrine signaling, these spatial relationships between *Bdnf* Ex1-expressing cells may be functionally relevant for BDNF-dependent structural and synaptic plasticity (Harward *et al*, 2016; Hedrick *et al*, 2016; Song *et al*, 2017).

We focused specifically on the piriform cortex because this region expresses high levels of BDNF, and is highly epileptogenic; hence, this brain region may be particularly responsive to ECS. The piriform cortex facilitates temporal seizure kindling and propagates generalized seizures that are induced by repeated electrical stimulations (Löscher and Ebert, 1996; Morimoto *et al*, 2004). In addition, the piriform cortex has been more recently linked to depression (Kohli *et al*, 2016; Sevelinges *et al*, 2011), and is functionally connected to to many regions relevant for MDD including the prefrontal cortex (Carmichael *et al*, 1994; Cohen *et al*, 2015; Gottfried and Zelano, 2011; Johnson *et al*, 2000; Kringelbach and Rolls, 2004; Plailly *et al*, 2008), amygdala (Carmichael *et al*, 1994; Johnson *et al*, 2000; Kajiwara *et al*, 2007; Kjelvik *et al*, 2012; Krettek and Price, 1977; Royet *et al*, 2003; Winston *et al*, 2005), subiculum/hippocampus (Kjelvik *et al*, 2012; Krettek and Price, 1977), and the hypothalamus (Price *et al*, 1991). Further supporting a potential involvement in the response to stress, the expression of proteins implicated in structural plasticity, such as PSA-NCAM and doublecortin, are decreased in piriform cortex in rodent models of chronic stress (Nacher *et al*, 2004).

We hypothesized that expression of *Bdnf* Ex1 is required for spine remodeling following ECS in *Bdnf* Ex1-expressing neurons in the piriform cortex. However, studying cellular morphology in *Bdnf* Ex1-expressing cells required a strategy to genetically label this population. Unfortunately, molecular genetic tools for selectively tagging populations of cells based on transcript-specific expression of *Bdnf* are limited (Maynard *et al*, 2016, 2017). We took advantage of the Cre-recombinase dependent on GFP (CRE-DOG) system, which we used in combination with GFP-expressing Bdnf-e1 mice. These mice report transcriptional activity from *Bdnf* promoter I while simultaneously disrupting functional *Bdnf* expression from this promoter. Since CRE-DOG requires GFP for scaffolding, we were limited in that we could only compare dendritic morphology in heterozygous mice that expressed BDNF produced from promoter I at one allele and homozygous mutant mice that fail to produce any BDNF from this promoter. It is possible that the ~50% reduction in promoter I-derived BDNF in heterozygous animals may have prevented the robust ECS-induced spine changes we previously observed in wild-type animals (Maynard *et al*, 2018). Conducting similar studies that would allow us to compare changes across wild-type, heterozygous and mutant mice would require development and optimization of novel tools. For example, we recently generated a *Bdnf* promoter IV tamoxifen-inducible Cre system to selectively label the cell population expression BDNF from promoter IV in the hippocampus (Hallock *et al*, 2019), and it is possible that a similar strategy could be used with *Bdnf* promoter I.

In this study, mutant Bdnf-e1 −/− mice showed no increase in spine density in *Bdnf* Ex1-expressing neurons following repeated ECS - indeed, we actually observed a decrease in the percentage of smaller spine populations (widths 0.2-0.4 μm). It should be noted that while decreases in spine density and populations of smaller spines are observed in Bdnf-e1 −/− Sham mice, these effects are non-significant, suggesting that repeated stimulation may make dendrites more vulnerable to spine impairments. Given that smaller spine populations are reduced following ECS, *Bdnf* Ex1 transcripts may play a crucial role in stabilizing and maintaining newly formed spines, which are smaller in diameter than mature spines. This is consistent with our previous findings showing that ECS administration in a neuroendocrine model of chronic stress significantly impacts spine diameter in the 0.2 μm to 0.4 μm range (Maynard *et al*, 2018). The most robust changes we observed were in the population of spines with widths >0.2 μm, indicating that spine maintenance was affected by ECS. In summary, our results demonstrate that individual *Bdnf* splice isoforms are differentially regulated in response to ECS, and that cells expressing these transcripts show unique spatial patterns of activation and recruitment in response to ECS. Specifically, *Bdnf* Ex1-expressing cells are preferentially recruited in response to ECS, which may be functionally relevant since promoter I-derived *Bdnf* is required for ECS-induced structural plasticity in *Bdnf* Ex1-expressing neurons.

## ACKNOWLEDGEMENTS

We thank members of the Neural Plasticity groups at LIBD for helpful comments and advice on the manuscript.

## AUTHOR CONTRIBUTIONS

AR: conceptualization of work, interpretation of data, writing the original draft, and editing of intellectual content. KRM: conceptualization and design of work, data analysis, interpretation of data, and editing of intellectual content. AK: conceptualization and design of work, data acquisition and analysis, interpretation of data. BP: conceptualization, software development, and contributed new analytic tools SR: data acquisition and analysis. MT: software development, data analysis, and data visualization. JWH: data acquisition and analysis. SCP: conceptualization, data acquisition, and analysis. AEJ: conceptualization, statistical analyses, software development, contributed new analytic tools, and resources. KM: conceptualization and design of work, editing of intellectual content, supervision, resources and funding acquisition.

## REFERENCES

Aid T, Kazantseva A, Piirsoo M, Palm K, Timmusk T (2007). Mouse and rat BDNF gene structure and expression revisited. J Neurosci Res 85: 525–535.

Alfarez DN, De Simoni A, Velzing EH, Bracey E, Joëls M, Edwards FA, et al (2009). Corticosterone reduces dendritic complexity in developing hippocampal CA1 neurons. Hippocampus 19: 828–836.

Arellano JI, Benavides-Piccione R, Defelipe J, Yuste R (2007). Ultrastructure of dendritic spines: correlation between synaptic and spine morphologies. Front Neurosci 1: 131–143.

Autry AE, Adachi M, Nosyreva E, Na ES, Los MF, Cheng P, et al (2011). NMDA receptor blockade at rest triggers rapid behavioural antidepressant responses. Nature 475: 91–95.

Autry AE, Monteggia LM (2012). Brain-derived neurotrophic factor and neuropsychiatric disorders. Pharmacol Rev 64: 238–258.

Azis IA, Hashioka S, Tsuchie K, Miyaoka T, Abdullah RA, Limoa E, et al (2019).Electroconvulsive shock restores the decreased coverage of brain blood vessels by astrocytic endfeet and ameliorates depressive-like behavior. J Affect Disord 257: 331–339.

Baj G, Leone E, Chao MV, Tongiorgi E (2011). Spatial segregation of BDNF transcripts enables BDNF to differentially shape distinct dendritic compartments. Proc Natl Acad Sci USA 108: 16813–16818.

Bohlen Und Halbach O von, Bohlen Und Halbach V von (2018). BDNF effects on dendritic spine morphology and hippocampal function. Cell Tissue Res 373: 729–741.

Carmichael ST, Clugnet MC, Price JL (1994). Central olfactory connections in the macaque monkey. J Comp Neurol 346: 403–434.

Chang AD, Vaidya PV, Retzbach EP, Chung SJ, Kim U, Baselice K, et al (2018). Narp Mediates Antidepressant-Like Effects of Electroconvulsive Seizures. Neuropsychopharmacology 43: 1088–1098.

Chen B, Dowlatshahi D, MacQueen GM, Wang JF, Young LT (2001). Increased hippocampal BDNF immunoreactivity in subjects treated with antidepressant medication. Biol Psychiatry 50: 260–265.

Chiaruttini C, Sonego M, Baj G, Simonato M, Tongiorgi E (2008). BDNF mRNA splice variants display activity-dependent targeting to distinct hippocampal laminae. Mol Cell Neurosci 37: 11–19.

Cohen Y, Wilson DA, Barkai E (2015). Differential modifications of synaptic weights during odor rule learning: dynamics of interaction between the piriform cortex with lower and higher brain areas. Cereb Cortex 25: 180–191.

Duman CH, Duman RS (2015). Spine synapse remodeling in the pathophysiology and treatment of depression. Neurosci Lett 601: 20–29.

Duman RS, Monteggia LM (2006). A neurotrophic model for stress-related mood disorders. Biol Psychiatry 59: 1116–1127.

Ekstrand JJ, Domroese ME, Johnson DM, Feig SL, Knodel SM, Behan M, et al (2001). A new subdivision of anterior piriform cortex and associated deep nucleus with novel features of interest for olfaction and epilepsy. J Comp Neurol 434: 289–307.

Goldwater DS, Pavlides C, Hunter RG, Bloss EB, Hof PR, McEwen BS, et al (2009). Structural and functional alterations to rat medial prefrontal cortex following chronic restraint stress and recovery. Neuroscience 164: 798–808.

Gottfried JA, Zelano C (2011). The value of identity: olfactory notes on orbitofrontal cortex function. Ann N Y Acad Sci 1239: 138–148.

Guilloux JP, Douillard-Guilloux G, Kota R, Wang X, Gardier AM, Martinowich K, et al (2012). Molecular evidence for BDNF- and GABA-related dysfunctions in the amygdala of female subjects with major depression. Mol Psychiatry 17: 1130–1142.

Hallock HL, Quillian HM, Mai Y, Maynard KR, Hill JL, Martinowich K (2019). Manipulation of a genetically and spatially defined sub-population of BDNF-expressing neurons potentiates learned fear and decreases hippocampal-prefrontal synchrony in mice. Neuropsychopharmacology 44: 2239–2246.

Hallock HL, Quillian HM, Maynard KR, Mai Y, Chen H-Y, Hamersky GR, et al (2020). Molecularly defined hippocampal inputs regulate population dynamics in the prelimbic cortex to suppress context fear memory retrieval. Biol Psychiatry 88: 554–565.

Harward SC, Hedrick NG, Hall CE, Parra-Bueno P, Milner TA, Pan E, et al (2016). Autocrine BDNF-TrkB signalling within a single dendritic spine. Nature 538: 99–103.

Hedrick NG, Harward SC, Hall CE, Murakoshi H, McNamara JO, Yasuda R (2016). Rho GTPase complementation underlies BDNF-dependent homo- and heterosynaptic plasticity. Nature 538: 104–108.

Hering H, Sheng M (2001). Dendritic spines: structure, dynamics and regulation. Nat Rev Neurosci 2: 880–888.

Hill JL, Hardy NF, Jimenez DV, Maynard KR, Kardian AS, Pollock CJ, et al (2016). Loss of promoter IV-driven BDNF expression impacts oscillatory activity during sleep, sensory information processing and fear regulation. Transl Psychiatry 6: e873.

Johnson DM, Illig KR, Behan M, Haberly LB (2000). New features of connectivity in piriform cortex visualized by intracellular injection of pyramidal cells suggest that “primary” olfactory cortex functions like “association” cortex in other sensory systems. J Neurosci 20: 6974–6982.

Kaastrup Müller H, Orlowski D, Reidies Bjarkam C, Wegener G, Elfving B (2015). Potential roles for Homer1 and Spinophilin in the preventive effect of electroconvulsive seizures on stress-induced CA3c dendritic retraction in the hippocampus. Eur Neuropsychopharmacol 25: 1324–1331.

Kajiwara R, Tominaga T, Takashima I (2007). Olfactory information converges in the amygdaloid cortex via the piriform and entorhinal cortices: observations in the guinea pig isolated whole-brain preparation. Eur J Neurosci 25: 3648–3658.

Kho KH, Vreeswijk MF van, Simpson S, Zwinderman AH (2003). A meta-analysis of electroconvulsive therapy efficacy in depression. J ECT 19: 139–147.

Kjelvik G, Evensmoen HR, Brezova V, Håberg AK (2012). The human brain representation of odor identification. J Neurophysiol 108: 645–657.

Kohli P, Soler ZM, Nguyen SA, Muus JS, Schlosser RJ (2016). The association between olfaction and depression: A systematic review. Chem Senses 41: 479–486.

Kraus C, Kadriu B, Lanzenberger R, Zarate CA, Kasper S (2019). Prognosis and improved outcomes in major depression: a review. Transl Psychiatry 9: 127.

Krettek JE, Price JL (1977). Projections from the amygdaloid complex and adjacent olfactory structures to the entorhinal cortex and to the subiculum in the rat and cat. J Comp Neurol 172: 723–752.

Kringelbach ML, Rolls ET (2004). The functional neuroanatomy of the human orbitofrontal cortex: evidence from neuroimaging and neuropsychology. Prog Neurobiol 72: 341–372.

Lee B-H, Kim Y-K (2010). The roles of BDNF in the pathophysiology of major depression and in antidepressant treatment. Psychiatry Investig 7: 231–235.

Löscher W, Ebert U (1996). The role of the piriform cortex in kindling. Prog Neurobiol 50: 427–481.

Martinowich K, Manji H, Lu B (2007). New insights into BDNF function in depression and anxiety. Nat Neurosci 10: 1089–1093.

Maynard KR, Hill JL, Calcaterra NE, Palko ME, Kardian A, Paredes D, et al (2016). Functional Role of BDNF Production from Unique Promoters in Aggression and Serotonin Signaling. Neuropsychopharmacology 41: 1943–1955.

Maynard KR, Hobbs JW, Rajpurohit SK, Martinowich K (2018). Electroconvulsive seizures influence dendritic spine morphology and BDNF expression in a neuroendocrine model of depression. Brain Stimulat 11: 856–859.

Maynard KR, Hobbs JW, Sukumar M, Kardian AS, Jimenez DV, Schloesser RJ, et al (2017). Bdnf mRNA splice variants differentially impact CA1 and CA3 dendrite complexity and spine morphology in the hippocampus. Brain Struct Funct 222: 3295–3307.

Maynard KR, Tippani M, Takahashi Y, Phan BN, Hyde TM, Jaffe AE, et al (2020). dotdotdot: an automated approach to quantify multiplex single molecule fluorescent in situ hybridization (smFISH) images in complex tissues. Nucleic Acids Res doi:10.1093/nar/gkaa312.

McAllan L, Maynard KR, Kardian AS, Stayton AS, Fox SL, Stephenson EJ, et al (2018). Disruption of brain-derived neurotrophic factor production from individual promoters generates distinct body composition phenotypes in mice. Am J Physiol Endocrinol Metab 315: E1168–E1184.

McKay MS, Zakzanis KK (2010). The impact of treatment on HPA axis activity in unipolar major depression. J Psychiatr Res 44: 183–192.

Morales-Medina JC, Sanchez F, Flores G, Dumont Y, Quirion R (2009). Morphological reorganization after repeated corticosterone administration in the hippocampus, nucleus accumbens and amygdala in the rat. J Chem Neuroanat 38: 266–272.

Morimoto K, Fahnestock M, Racine RJ (2004). Kindling and status epilepticus models of epilepsy: rewiring the brain. Prog Neurobiol 73: 1–60.

Nacher J, Pham K, Gil-Fernandez V, McEwen BS (2004). Chronic restraint stress and chronic corticosterone treatment modulate differentially the expression of molecules related to structural plasticity in the adult rat piriform cortex. Neuroscience 126: 503–509.

Nibuya M, Morinobu S, Duman RS (1995). Regulation of BDNF and trkB mRNA in rat brain by chronic electroconvulsive seizure and antidepressant drug treatments. J Neurosci 15: 7539–7547.

Pagnin D, Queiroz V de, Pini S, Cassano GB (2004). Efficacy of ECT in depression: a meta-analytic review. J ECT 20: 13–20.

Pattabiraman PP, Tropea D, Chiaruttini C, Tongiorgi E, Cattaneo A, Domenici L (2005). Neuronal activity regulates the developmental expression and subcellular localization of cortical BDNF mRNA isoforms in vivo. Mol Cell Neurosci 28: 556–570.

Plailly J, Howard JD, Gitelman DR, Gottfried JA (2008). Attention to odor modulates thalamocortical connectivity in the human brain. J Neurosci 28: 5257–5267.

Price JL, Slotnick BM, Revial MF (1991). Olfactory projections to the hypothalamus. J Comp Neurol 306: 447–461.

Radley JJ, Rocher AB, Rodriguez A, Ehlenberger DB, Dammann M, McEwen BS, et al (2008). Repeated stress alters dendritic spine morphology in the rat medial prefrontal cortex. J Comp Neurol 507: 1141–1150.

Royet J-P, Plailly J, Delon-Martin C, Kareken DA, Segebarth C (2003). fMRI of emotional responses to odors: influence of hedonic valence and judgment, handedness, and gender. Neuroimage 20: 713–728.

Russo-Neustadt AA, Beard RC, Huang YM, Cotman CW (2000). Physical activity and antidepressant treatment potentiate the expression of specific brain-derived neurotrophic factor transcripts in the rat hippocampus. Neuroscience 101: 305–312.

Sathanoori M, Dias BG, Nair AR, Banerjee SB, Tole S, Vaidya VA (2004). Differential regulation of multiple brain-derived neurotrophic factor transcripts in the postnatal and adult rat hippocampus during development, and in response to kainate administration. Brain Res Mol Brain Res 130: 170–177.

Savell KE, Bach SV, Zipperly ME, Revanna JS, Goska NA, Tuscher JJ, et al (2019). A Neuron-Optimized CRISPR/dCas9 Activation System for Robust and Specific Gene Regulation. Eneuro 6:.

Schloesser RJ, Orvoen S, Jimenez DV, Hardy NF, Maynard KR, Sukumar M, et al (2015). Antidepressant-like Effects of Electroconvulsive Seizures Require Adult Neurogenesis in a Neuroendocrine Model of Depression. Brain Stimulat 8: 862–867.

Schmidt HD, Shelton RC, Duman RS (2011). Functional biomarkers of depression: diagnosis, treatment, and pathophysiology. Neuropsychopharmacology 36: 2375–2394.

Sevelinges Y, Mouly A-M, Raineki C, Moriceau S, Forest C, Sullivan RM (2011). Adult depression-like behavior, amygdala and olfactory cortex functions are restored by odor previously paired with shock during infant’s sensitive period attachment learning. Dev Cogn Neurosci 1: 77–87.

Smith MA, Makino S, Kvetnansky R, Post RM (1995). Stress and glucocorticoids affect the expression of brain-derived neurotrophic factor and neurotrophin-3 mRNAs in the hippocampus. J Neurosci 15: 1768–1777.

Song M, Martinowich K, Lee FS (2017). BDNF at the synapse: why location matters. Mol Psychiatry 22: 1370–1375.

Soudry Y, Lemogne C, Malinvaud D, Consoli SM, Bonfils P (2011). Olfactory system and emotion: common substrates. Eur Ann Otorhinolaryngol Head Neck Dis 128: 18–23.

Tang JCY, Rudolph S, Dhande OS, Abraira VE, Choi S, Lapan SW, et al (2015). Cell type-specific manipulation with GFP-dependent Cre recombinase. Nat Neurosci 18: 1334–1341.

Timmusk T, Palm K, Metsis M, Reintam T, Paalme V, Saarma M, et al (1993). Multiple promoters direct tissue-specific expression of the rat BDNF gene. Neuron 10: 475–489.

UK ECT Review Group (2003). Efficacy and safety of electroconvulsive therapy in depressive disorders: a systematic review and meta-analysis. Lancet 361: 799–808.

Winston JS, Gottfried JA, Kilner JM, Dolan RJ (2005). Integrated neural representations of odor intensity and affective valence in human amygdala. J Neurosci 25: 8903–8907.

Yuuki N, Ida I, Oshima A, Kumano H, Takahashi K, Fukuda M, et al (2005). HPA axis normalization, estimated by DEX/CRH test, but less alteration on cerebral glucose metabolism in depressed patients receiving ECT after medication treatment failures. Acta Psychiatr Scand 112: 257–265.

